# Sleep spindle state-dependent motor cortical plasticity induction by EEG-triggered ripple burst TMS

**DOI:** 10.64898/2026.09.01.748506

**Authors:** Zhijian Zhao, Yeyun Lu, Friederike Breuer, Til Ole Bergmann, Ulf Ziemann

## Abstract

**Background:** Sleep spindles are fundamental for plasticity and memory consolidation. Here we sought to target different spindle states in sleeping healthy participants with real-time EEG-burst repetitive transcranial magnetic stimulation (rTMS) at hippocampal ripple frequency, and test the spindle state-dependent induction of corticospinal and sensorimotor cortical plasticity. We hypothesized that the spindle-trough is a particularly critical state for plasticity induction because hippocampal ripples are naturally nested in the spindle-trough, reflecting replay of memory traces and facilitating memory consolidation.

**Methods:** Fourteen participants underwent four experimental nights, in which rTMS was applied either at the spindle-trough, spindle-peak, spindle random phase or during spindle-free epochs. Readouts of plasticity were changes in resting-state EEG (rsEEG) power, motor evoked potential (MEP) amplitude, local mean field amplitude (LMFA, for the N45 and P60 potential components), immediate response slope (IRS) and TMS-related time frequency response (TFR), tested 10 and 30 min after the end of the rTMS interventions upon awakening, and compared to pre-sleep baseline.

**Results:** Spindle-trough rTMS resulted in pre- to post-sleep decreases of rsEEG beta-band power, MEP amplitude, P60-LMFA, IRS and TFR in the alpha-band, and an increase in the N45-LMFA. None of the other spindle state-dependent rTMS interventions resulted in consistent plastic changes.

**Conclusions:** Targeting the spindle-trough with ripple-burst rTMS stands out in consistently leading to long-term depression-like changes across a broad array of corticospinal and sensorimotor cortical excitability readouts. This opens the intriguing opportunity of targeted manipulation of human sleep physiology for improving specific behavioral processes, such as memory consolidation.

## 1 Introduction

Sleep spindles are brief sigma-band (11-15 Hz) oscillations generated in thalamocortical circuits during non-rapid eye movement (NREM) sleep (De Gennaro & Ferrara, 2003; Jankel & Niedermeyer, 1985). They occur within a broader temporal structure of NREM activity involving slow oscillations (SO, <1 Hz) and hippocampal sharp-wave ripples (>80 Hz). Spindles can be nested within this slower rhythm, and hippocampal ripple-related activity preferentially occurs at specific phases of the spindle cycle (Staresina et al., 2015, 2023). Given the strong Ca^+^ influx into cortical neurons associated with their occurrence (Rosanova & Ulrich, 2005), spindles have been proposed as transient windows for plasticity, with rhythmic thalamocortical events creating temporally structured conditions for cortical processing (Bergmann & Born, 2018; Fernandez & Lüthi, 2020). External stimulation delivered at different spindle phases may therefore produce different physiological effects.

Real-time EEG-triggered transcranial magnetic stimulation (TMS) allows stimulation to be delivered at specific phases of ongoing oscillatory activity (Bergmann et al., 2019; Schaworonkow et al., 2018; Stefanou et al., 2019; Zrenner et al., 2018, 2026). During wakefulness, the amplitude of corticospinal responses elicited by TMS of the primary motor cortex (M1), as indexed by motor evoked potentials (MEPs), varies with the phase and power of the sensorimotor mu-alpha rhythm, with the trough and ascending phase being a state of high corticospinal excitability, and the peak and descending phase reflecting low corticospinal excitability (Hussain et al., 2019; Wischnewski et al., 2022; Zrenner et al., 2018, 2023). Repetitive TMS targeting specific phases of the mu-rhythm has demonstrated induction of differential corticospinal plasticity. High-frequency TMS triplets targeted to the mu trough have been reported to induce a long-term potentiation (LTP)-like increase in corticospinal excitability (Baur et al., 2022; Zrenner et al., 2018), whereas low-frequency repetitive TMS timed to the mu peak induced a long-term depression (LTD)-like decrease (Baur et al., 2020). Phase-dependent stimulation effects are not limited to the motor system and mu-rhythm. Prefrontal stimulation targeting different phases of the local theta-rhythm differentially modulated subsequent TMS-EEG responses and working-memory performance (Gordon et al., 2022).

Direct evidence on how corticospinal excitability varies across sleep spindle phase remains limited. In a recent single-pulse TMS study of centroparietal spindles (12-15 Hz), spindle-phase-triggered TMS over M1 showed lower MEP amplitude during spindles, driven by particular suppression during the falling flank, compared to any other targeted phase, spindle-free periods, or immediate post-spindle refractory periods (Hassan et al., 2025). This finding was principally replicated in an independent sample showing spindle-locked suppression, nominally strongest during its falling flank (Breuer et al., bioRxiv 2026). Findings provide evidence that corticospinal excitability is phase-modulated during spindles and that specific spindle phases can be reliably targeted in real time. However, the differential effects of repetitive TMS delivered at selected fast-spindle phases on plasticity induction have not yet been tested. This is a fundamental neurophysiological question, as SO-spindle-ripple events are assumed to provide a window of enhanced synaptic plasticity that facilitates the redistribution of memory representations across large-scale networks (Bergmann & Born, 2018; Klinzing et al., 2019; Staresina, 2024), and the targeted modulation of this process might eventually serve the improvement of memory processes.

In the present study, we tested the effects of repetitive TMS targeting different spindle phases on the induction of corticospinal excitability. Repeated TMS triplet bursts in the hippocampal ripple frequency range were delivered in four separate nocturnal nap sessions at the peak, trough, or at random phases of real-time EEG-detected fast spindles in M1, or in spindle-free periods. We hypothesized that corticospinal plasticity would vary between the four conditions, with stimulation targeting the spindle trough revealing the most pronounced effects, given that (i) this is the natural spindle phase in which hippocampal ripples are predominantly nested (Staresina et al., 2015), and (ii) previous work using mu-alpha phase-triggered TMS of M1 during wakefulness has found this phase to show the most consistent phase-dependent plasticity effects.

## 2 Methods

### 2.1 Participants

Of 38 screened healthy participants, 14 (8 females, 6 males; age = 27.0 ± 2.4 years, range: 23-31) contributed to the final dataset. The other 24 screened individuals were not included because they did not meet the inclusion criteria, in particular their ability to sleep in a supine position for at least one hour during the adaptation night, or the presence of a reliable TMS motor hotspot. Exclusion criteria included present or past neurological or psychiatric disorders, pregnancy, irregular sleep schedules (e.g., due to shift work), excessive consumption of caffeinated beverages, regular drug use or history of drug abuse, and contraindications to TMS (Rossi et al., 2021). Inclusion required (1) the ability to sleep supine for at least one hour under the experimental setup without strong head movements and (2) the presence of a reliable TMS motor hotspot, defined as a scalp location at which TMS elicited MEPs of at least 0.5 mV peak-to-peak amplitude in the right first dorsal interosseous (FDI) or abductor pollicis brevis (APB) muscle during screening in wakefulness and while wearing an EEG cap. All participants gave written informed consent before participation. The study conformed to the last version of the Declaration of Helsinki and was approved by the ethics committee of the Medical Faculty of the University of Tübingen (project: 349/2018BO2).

The final dataset comprised 14 spindle peak, 14 spindle trough, 13 spindle random-phase and 14 spindle-free conditions (see below for definition of experimental conditions), since one participant withdrew voluntarily before completing the random-phase night. An a priori minimum number of 120 stimulation events was required for session inclusion.

A sensitivity analysis was performed in G*Power using a four-level repeated-measures ANOVA as an approximation of the within-subject condition effect. With 14 participants, α = 0.05, and 80% power, the minimum detectable effect was Cohen’s f= 0.365, assuming a correlation of 0.50 among repeated measures and a non-sphericity correction of ε= 0.75.

### 2.2 Experimental design and general procedure

Participants completed one adaptation nocturnal-nap session followed by four nocturnal experimental nights in the Brain Networks & Plasticity Lab at the University of Tübingen. The adaptation session familiarized participants with the sleep environment, including sleeping in a supine position with the head fixed by a customized pillow with a U-shaped groove, while their EEG, electrooculography (EOG) and electromyogram (EMG) were recorded. The EEG data provided information on individual spindle frequency for optimization of real-time spindle detection during the nocturnal experimental sessions. Each experimental session included a nocturnal nap during the first part of the participant’s habitual sleep period. Each nocturnal experimental session was assigned to one TMS condition: spindle peak, spindle trough, spindle random phase, or spindle-free N2/N3 sleep (Fig. 1B). The order of the four experimental sessions was pseudorandomized and balanced across participants, and experimental sessions were separated by at least 3 days to avoid carryover effects. Participants were blinded to the stimulation condition, whereas experimenters were aware of the assigned condition because the real-time stimulation protocol had to be set up before each intervention.

**Figure 1.**
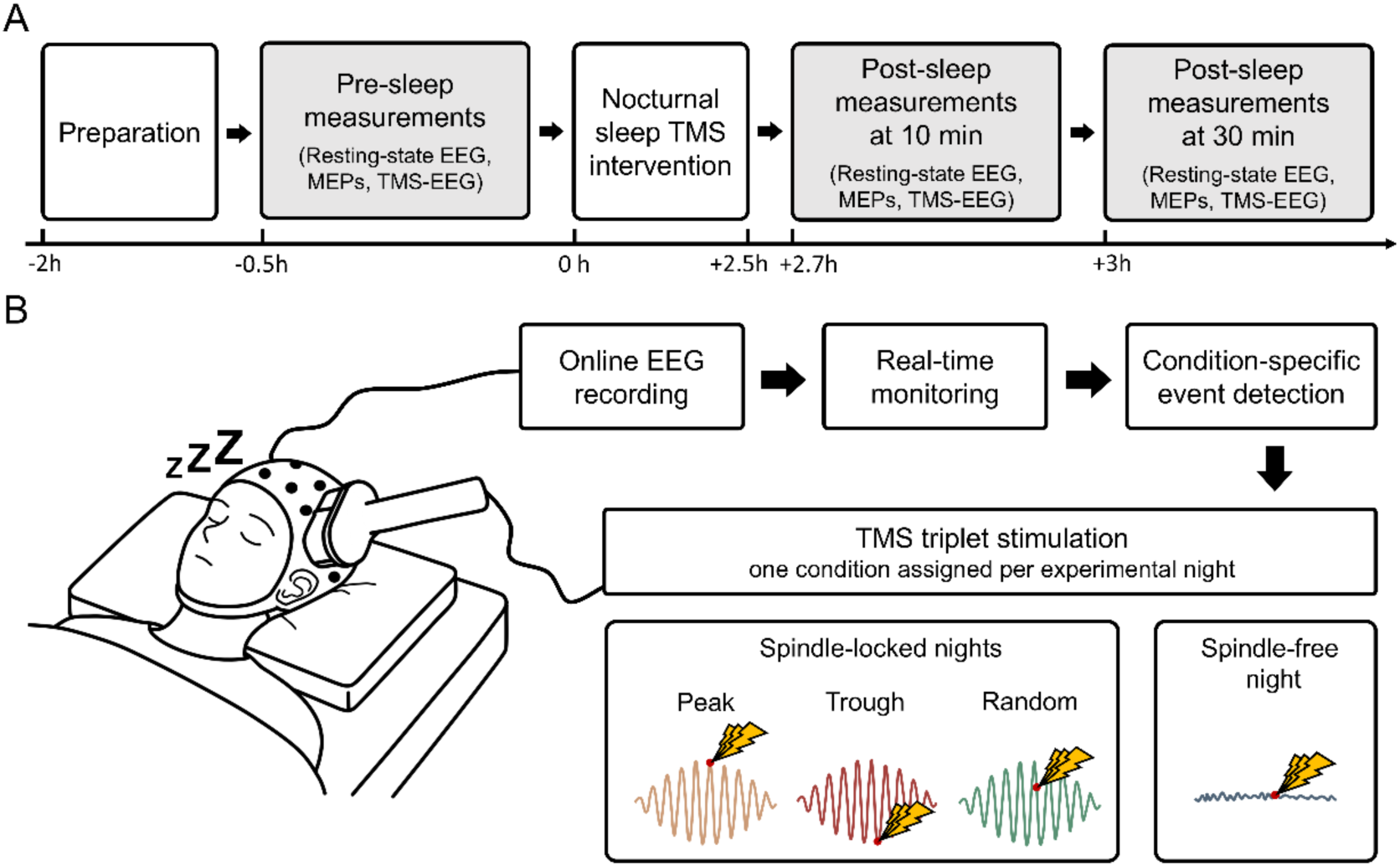
Experimental session timeline and real-time EEG triggered TMS triplet stimulation. (A) Timeline for a single experimental session. Preparation was followed by pre-sleep measurements, the sleep TMS intervention, and post-sleep measurements at 10 and 30 min. (B) Real-time EEG triggered TMS triplet stimulation during the sleep intervention. Each experimental session was assigned to one stimulation condition. In spindle-locked sessions, TMS triplets were delivered at the peak, trough, or at random phase of detected fast spindles. In spindle-free sessions, triplets were delivered outside spindle periods.

On each experimental session, participants arrived approximately 2 h before their usual bedtime and were prepared for sleep EEG/EOG/EMG recordings, TMS neuronavigation, motor-hotspot localization, automated resting motor threshold (RMT) estimation, and pre-sleep measurements of eyes-open resting-state EEG (rs-EEG), MEP, and TMS-EEG (Fig. 1A). Participants then slept in supine position in a soundproof and electromagnetically shielded sleeping cabin (Desone, Germany), with earplugs, and the head stabilized using a dedicated fixation pillow. The TMS coil was held in position by a mechanical arm. During sleep, TMS bursts (triplets at 85 Hz) were real-time EEG triggered during NREM stages N2 or N3 when the criteria for the assigned experimental condition were met. Sleep stages were evaluated online by a trained experimenter in consecutive 30-s epochs using the continuous EEG, EOG, and EMG recordings, following the standardized American Academy of Sleep Medicine (AASM) criteria (Berry et al., 2020). Outcome measures (rs-EEG, MEPs, and TMS-EEG) were recorded again 10 and 30 min after awakening (for details, please see below).

### 2.3 EEG and sleep recordings

Sleep recordings included 64-channel EEG, EOG and surface EMG. EEG was acquired using a TMS-compatible NeurOne Tesla system with a 24-bit battery-powered amplifier (NeurOne Tesla with Digital-Out Option, Bittium, Finland). Signals were recorded with sintered Ag/AgCl electrodes mounted in a 64-channel TMS-compatible cap (Multitrodes-TMS, EasyCap; channels Fp1, Fp2, F3, F4, C3, C4, P3, P4, O1, O2, F7, F8, T7, T8, P7, P8, AFz, Fz, FCz, Pz, FC1, FC2, CP1, CP2, FC5, FC6, CP5, CP6, FT9, FT10, F1, F2, C1, C2, P1, P2, AF3, AF4, FC3, FC4, CP3, CP4, PO3, PO4, F5, F6, C5, C6, P5, P6, AF7, AF8, FT7, FT8, TP7, TP8, PO7, PO8, Fpz, Cz, POz, Oz, M1, and M2; reference: CPz). Electrode impedances were kept below 5 kΩ throughout recording. EOG electrodes were placed approximately 1 cm above the outer canthus of one eye and 1 cm below the outer canthus of the contralateral eye and referenced to contralateral mastoids, following the AASM scoring recommendations. Surface EMG was recorded from the FDI and APB muscles of the right hand, using disposable gel electrodes in a belly-tendon montage. All signals were acquired in DC mode at 5 kHz, with a 1250-Hz hardware low-pass filter applied during acquisition. Sleep was monitored throughout the experiment using EEG, EOG. The EMG of the chin and the right hand was used to monitor muscle activity during sleep, and infrared video monitoring was used in parallel to identify gross body movements, posture changes, or visible awakenings.

### 2.4 TMS and neuronavigation

TMS was delivered over the left M1 hand area. Before sleep, the motor hotspot was identified as the scalp location where slightly suprathreshold TMS elicited the most consistent MEPs in the right FDI or APB muscle (Rossini et al., 2015). The right FDI was selected as the primary target muscle. If a stable FDI hotspot and reproducible MEPs could not be obtained with EEG cap in place and under the sleep-experimental setup, the right APB was used instead. The selected target muscle was kept the same across all experimental sessions and measurement sessions within a given participant. The hotspot and coil orientation were saved and maintained using MR-template-based frameless stereotactic neuronavigation (Localite GmbH, Germany). The BEST toolbox was used to identify the motor hotspot and estimate RMT using an automated closed-loop staircasing procedure (Hassan et al., 2022). TMS was delivered using a compact, cooled Cool-B35 HO butterfly coil connected to a MagPro X100 stimulator (MagVenture, Denmark) over the M1 hotspot in the supine sleep setup. Biphasic current was induced in M1, with the second phase in posterior-to-anterior direction. Awake single-pulse MEP and TMS-EEG measurements were performed at 120% RMT or 100% maximum stimulator output, whichever was lower. The realized stimulation intensity for awake measurements was 117.43 ± 4.44% RMT. The inter-trial interval (ITI) between consecutive single TMS pulses ranged from 2-3 s for MEP, and 6-7 s for TMS-EEG. To reduce TMS-click-related auditory evoked potentials in the TMS-EEG recordings, participants wore in-ear headphones delivering continuous TMS-masking noise generated by the TAAC toolbox (Russo et al., 2022), with individualized volume intensity calibration. No masking noise was played during sleep.

During N2/N3 sleep, the intervention consisted of low-intensity triplet TMS bursts at 90% RMT. Each stimulation event included three pulses delivered at 85 Hz. A refractory interval of 6 s was enforced after each stimulation event, to avoid unintended induction of corticospinal plasticity by rTMS per se, which may occur at rTMS frequencies of > 0.5 Hz but are not to be expected with the frequency of < 0.16 Hz we have applied (Ziemann et al., 2008). The interval between successive events was therefore determined by both target-state occurrence and this imposed refractory interval.

### 2.5 Real-time EEG-TMS

The Real-Time Spindle Detector (RTSD) was used to detect sleep spindles and their instantaneous phase in real time, as described previously (Hassan et al., 2022, 2025), implemented into a customized earlier version of the bossdevice (sync2brain GmbH, Tübingen, Germany). A bipolar C3-M2 montage was used to extract centroparietal fast spindle activity over the left sensorimotor region. EEG recorded during the adaptation night was analyzed with YASA (https://github.com/raphaelvallat/yasa) to estimate each participant’s individual fast-spindle peak frequency and sigma-band root mean square (RMS) threshold for the RTSD (Hassan et al., 2022; Vallat & Walker, 2021). During the nocturnal nap in each experimental session, sleep and signal quality were monitored online and sleep stages were scored visually according to AASM criteria. Formal scoring in fixed 30-s epochs was not used for stimulation control, as changes in sleep stability, arousals, and movements had to be detected on a shorter time scale. Real-time EEG-guided TMS was manually initiated after participants had entered stable NREM stage N2 sleep. TMS triplets were delivered during NREM stage N2 and N3, and were manually paused during wakefulness, N1 sleep, REM sleep, awakenings, major movements, or unstable sleep.

### 2.6 Validation of spindle-state and phase targeting

Offline, EEG data were analyzed immediately prior to the TMS pulse to verify that stimulation was delivered during the intended spindle-defined states and phases (spindle-trough, spindle-peak, spindle random phase, spindle-free).

Spindle-state at the time of the TMS triplets was evaluated offline using the C3-M2 derivation, matching the channel derivation used for real-time spindle detection. The signal was band-pass filtered in the fast-spindle range (12-15 Hz). Because EEG at and immediately after the TMS triplet was contaminated by stimulation-related artifacts, instantaneous phase was not estimated at the trigger itself. Instead, for each stimulation event, the dominant spindle frequency was estimated from the pre-trigger signal, and phase was extracted one spindle cycle before the first TMS pulse. Phase distributions and spindle-detection accuracy were visualized using polar histograms (Fig. 2A). Deviations from uniformity were assessed using Rayleigh’s test. For peak and trough stimulation, clustering toward the intended target phase was additionally assessed using the V-test. The V-test was not applied to random-phase or spindle-free stimulation, because these conditions did not have a predefined target phase. For each stimulation condition, sigma-filtered C3-M2 waveforms were averaged from -0.5 to - 0.005 s before the first TMS pulse of the triplet to visualize spindle activity and the targeted phase while avoiding the artifact-contaminated stimulation period (Fig. 2B).

**Figure 2.**
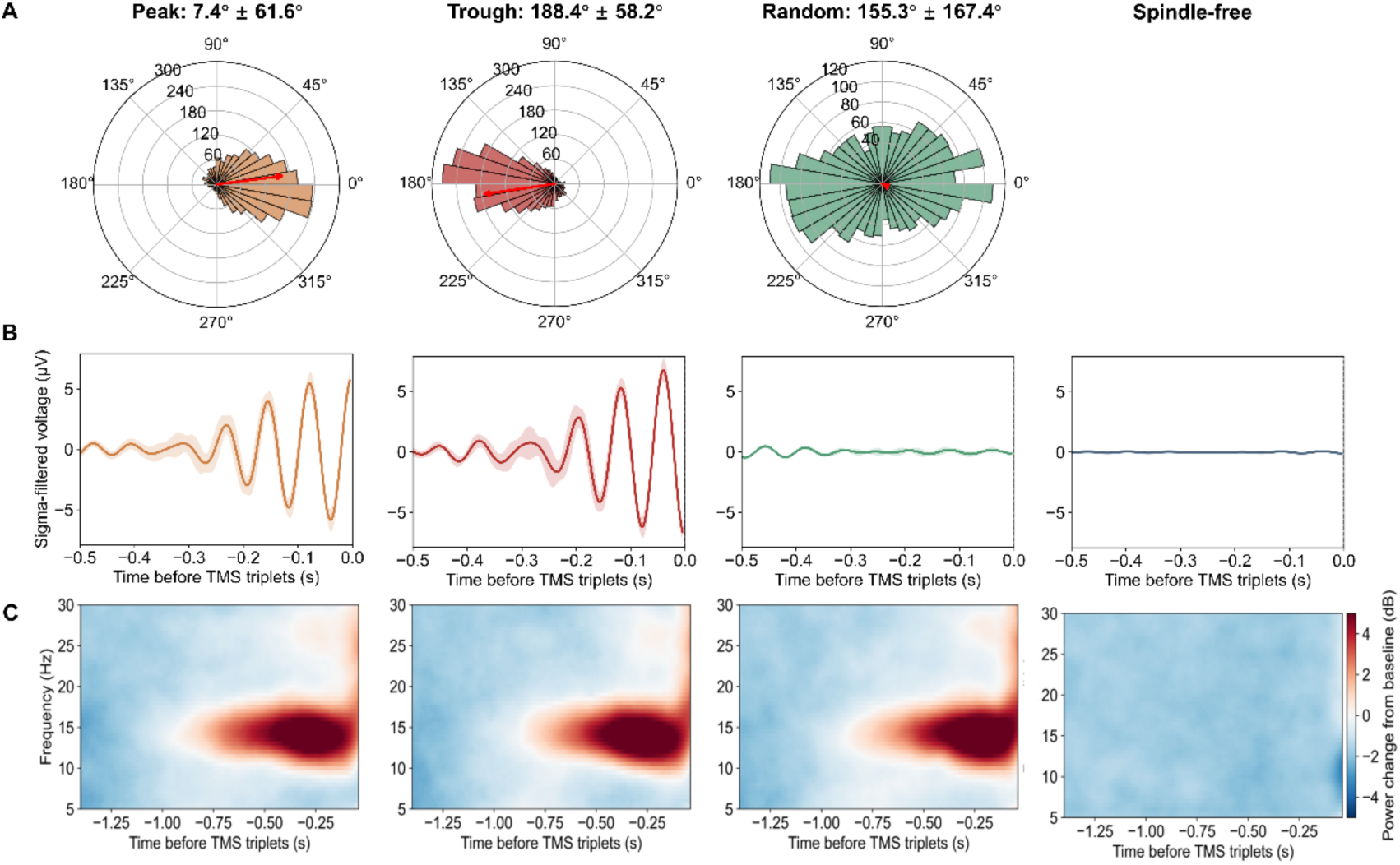
Offline analysis of spindle-state and phase targeting. (A) Polar histograms show spindle phase estimates for peak (0°), trough (180°), and random-phase stimulation, indicating the phase at the time of the first TMS pulse of the TMS triplet. 90° is the mid-descending phase, and 270° the mid-ascending phase. Rings represent the number of observations per phase bin. Red arrows indicate the circular mean phase, with arrow length representing the mean resultant length (*r*). Circular mean phase ± circular SD is reported for each condition above the polar plots. (B) Sigma-filtered C3-M2 waveforms show condition-averaged pre-trigger activity. The time of the first TMS pulse in the TMS triplet is a 0 s. Peak and trough stimulation showed the expected phase alignment, whereas random-phase stimulation showed reduced phase-locked activity after averaging across random phases. Spindle-free stimulation showed little organized sigma activity. Lines indicate grand averages across participants, and shadings indicate ± SEM. (C) Time-frequency representations show highly similar pre-trigger sigma activity in the trough, peak and random spindle-phase conditions but not in the spindle-free condition.

### 2.7 Pre- and post-sleep analyses

#### 2.7.1 Resting-state EEG preprocessing and analysis

A 5-min eyes-open rs-EEG recording was preprocessed offline using a semi-manual pipeline in Python/MNE (Gramfort, 2013). Continuous recordings were imported and visually inspected to verify channel layout, event information, recording quality, and the presence of gross artifacts. Data were band-pass filtered from 0.5 to 40 Hz and down-sampled to 500 Hz for spectral analyses. During visual inspection, channels with persistent noise, unstable contact, flat signal, excessive drift, or non-physiological activity were marked as bad. These channels were excluded before independent component analysis (ICA) estimation. ICA was applied to the cleaned continuous data to identify stereotypical ocular, cardiac, muscle, line-noise, and other non-brain signal components based on their time course, scalp topography, power spectrum, and automated component classification when available (Delorme & Makeig, 2004; Hyvärinen & Oja, 2000). After applying the ICA weights to the analysis dataset, artifact-related components were manually rejected. The removed EEG channels were then reconstructed using spherical interpolation based on the original channel locations, and the data were then re-referenced to the common average. This reference scheme was applied consistently across participants and sessions. After ICA correction, the data were visually inspected again, and remaining artifacts were removed if necessary. The final cleaned rs-EEG was saved for spectral and statistical analyses.

For the main rs-EEG sensorimotor analysis, spectral features were extracted from the cleaned EEG at pre-sleep (Pre), post-sleep 10 min (Post10), and post-sleep 30 min (Post30).

The primary sensor-level analyses used a predefined left-hemispheric sensorimotor region of interest (ROI) comprising FC1, FC3, C1, C3, CP1, and CP3. Power spectral density was estimated using Welch’s method. Analyses focused on mu/alpha (8-14 Hz), low beta (15-25 Hz), and high beta (25-30 Hz) bands.

#### 2.7.2 EMG analysis

MEP amplitudes were quantified from the EMG channel corresponding to the participant-specific target hand muscle. The right FDI muscle was used in 12 participants, whereas the right APB muscle was used in 2 participants for whom APB yielded more reliable MEPs.

Each MEP measurement block consisted of 50 single-pulse TMS trials. EMG data were epoched from -50 to 150 ms relative to the TMS pulse and baseline-corrected using the -50 to -10 ms prestimulus interval. Prestimulus background EMG activity was assessed in the same -50 to -10 ms window. Trials were excluded when prestimulus activity indicated pre-activation of the target muscle, defined as a rolling root-mean-square (RMS) exceeding 50 µV together with at least five consecutive samples exceeding a robust z-score threshold of 3. Robust z-scores were calculated relative to the prestimulus EMG distribution within each measurement block. For each retained trial, MEP amplitude was calculated as the peak-to-peak difference between the maximum and minimum EMG deflection within the 20-60 ms post-stimulation window. Remaining peak-to-peak outliers were excluded using a 1.5 * interquartile range rule (IQR) criterion (Stephan et al., 2018). Across all measurement blocks, 3.12 ± 2.70 trials (6.2% of trials) were excluded. No significant interaction between experimental condition and measurement time point was observed for the proportion of excluded trials (p = 0.833). Trial-level MEP amplitudes were log-transformed and subsequently averaged within each participant, stimulation condition, and time point for statistical analyses.

#### 2.7.3 TMS-EEG preprocessing and analysis

TMS-EEG recordings were preprocessed in MATLAB using EEGLAB (Delorme & Makeig, 2004; Hyvärinen & Oja, 2000) and TESA (Rogasch et al., 2017). Each TMS-EEG measurement block consisted of 100 single-pulse trials. Data were epoched from - 1500 to 1500 ms around each pulse, baseline corrected over -500 to -50 ms, and ultimately epochs were retained from -500 to 800 ms. The initial pulse artifact from - 2 to 4 ms was removed and reconstructed by cubic interpolation. Bad channels and trials were identified using amplitude-, robust-z-score-, RMS-based criteria, with rejection capped at 5% of channels and 10% of trials per recording. Data were resampled to 1000 Hz before decomposition. For ICA, the data rank was compressed to a maximum of 35 principal components and decomposed with symmetric FastICA using a *tanh* nonlinearity (Hyvarinen and Oja, 2000). Automated TESA component features were used to identify only clearly artifactual components, including early TMS-related muscle activity, blinks, movement-related activity, and electrode- or line-noise-like activity. On average, 6.18 ± 3.07 independent components were removed per recording. Residual channel-specific noise was attenuated with the source-utilized noise-discarding algorithm (SOUND, λ = 0.03, 10 iterations, Mutanen et al., 2018), using the standard-sensor-coordinates-based lead field generated by TESA. Channels removed during bad-channel screening were reconstructed during this step. Residual early TMS-evoked muscle activity was then reduced with signal-space-projection-source-informed-reconstruction (SSP-SIR, Mutanen et al., 2016), following source-based TMS-EEG artifact-reduction principles. A second pulse-artifact segment from -3 to 10 ms was removed and interpolated. Finally, data were filtered from 1 to 80 Hz, baseline corrected again over -500 to -50 ms.

TMS-evoked potential (TEP) analysis. TEPs were obtained by averaging the retained trials within each participant, experimental condition, and time point (Pre, Post10, Post30). For evoked-response analyses, each channel was additionally baseline corrected over -100 to -10 ms. Global mean field amplitude (GMFA) was inspected as a recording-level quality-control measure. The primary inferential local measure was the local mean field amplitude (LMFA) within the predefined left-motor ROI comprising FC1, FC3, C1, C3, CP1, and CP3 (Schoisswohl et al., 2024). At each sampled latency, LMFA was calculated as the spatial standard deviation of the voltages (Lehmann & Skrandies, 1980) across the six ROI channels. Analyses of predefined TEP component amplitudes were restricted to the first 80 ms after TMS onset to focus on early cortical responses and limit contribution from later auditory and somatosensory evoked activity (Biabani et al., 2019; Casarotto et al., 2013; Conde et al., 2019; Gordon et al., 2021). LMFA waveforms were visualized, and mean amplitudes were analyzed within predefined component windows, as follows: N15/early (10-25 ms), P30 (25-35 ms), N45 (40-50 ms), and P60 (50-75 ms). Post-intervention change scores were calculated separately as Post10-Pre and Post30-Pre.

As an exploratory measure of the early TEP response, evoked voltage signals were baseline-corrected from -100 to -10 ms. Baseline-corrected trials were first averaged at each electrode to obtain the evoked TEP. The resulting voltage waveforms were then averaged spatially across the six electrodes of the left sensorimotor ROI. The first temporal derivative was then estimated using finite differences, and its signed derivative was averaged over 10-25 ms after the TMS pulse. This yielded one immediate response slope (IRS) for each participant, intervention condition, and time point. The IRS was quantified using an adapted immediate response slope approach (Casarotto et al., 2013), with a fixed latency window applied to the trial-averaged TEP. For each time window and post-intervention assessment, differences among the four stimulation conditions (trough, peak, random, spindle-free) were tested using a repeated-measures spatial cluster-permutation test (Maris & Oostenveld, 2007), as described in Section 2.8.

For time-frequency representation (TFR) analysis, single-trial TMS-EEG signals were demeaned using the -400 to -100 ms interval and then averaged across the six ROI channels. Time-frequency power was estimated from 4 to 45 Hz in 1-Hz steps using Morlet wavelets. The number of wavelet cycles was set to frequency divided by two and constrained to a minimum of 3 and a maximum of 10 cycles. Calculations used Fast Fourier Transform (FFT) convolution and temporal decimation by a factor of four, resulting in TFR outputs with an effective temporal sampling rate of 250 Hz. Single-trial power was averaged across trials and converted to decibels relative to the -400 to - 100 ms baseline, defined as log10(power/baseline power). Post10-Pre and Post30-Pre TFR changes were calculated separately for each participant and condition.

### 2.8 Statistical analyses

Change-score analyses were performed for rs-EEG band power, log10-transformed MEP amplitude and LMFA at each timepoint of Post10 and Post30:

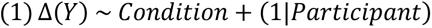

Here, Y denotes the analyzed outcome. The model included a fixed effect of condition and a random intercept for participant. Because Post10 and Post30 were modeled separately, the model did not include post-intervention time or an interaction between condition and post-invention time.

The overall effect of condition was evaluated using a joint Wald test. For each outcome family and post-intervention assessment, omnibus p values were adjusted using the Benjamini-Hochberg false discovery rate (FDR) procedure. Pairwise contrasts were performed only when the omnibus condition effect remained significant after FDR correction. All six pairwise contrasts among the four stimulation conditions (trough vs peak, trough vs random, trough vs spindle-free, peak vs random, peak vs spindle-free, and random vs spindle-free) were then tested. FDR correction was then applied across the six p values within the corresponding outcome family and post-intervention assessment.

Within each condition, participant-level Post10-Pre and Post30-Pre change scores were calculated against zero separately using two-sided one-sample t-tests. For each outcome family and assessment, FDR correction was applied across the p values from the corresponding condition-specific tests.

TMS-evoked time-frequency changes were evaluated separately at Post10 and Post30 using four-condition repeated-measures cluster-based permutation tests across 4-45 Hz and 0-300 ms. Clusters were defined using time-frequency adjacency and a cluster-forming threshold corresponding to p < 0.05. Cluster significance was estimated using 4096 permutations. For the omnibus condition tests, cluster-level p values were FDR corrected across the two post-intervention assessments. If the FDR-corrected omnibus condition effect was significant at a given assessment, all six pairwise condition contrasts were subsequently performed, with FDR correction applied across the six contrasts. FDR-adjusted p values are reported as p_corrected_, and statistical significance was set at α = 0.05.

## 3 Results

Results are organized to distinguish two levels of evidence: target engagement during the sleep intervention and pre-to-post changes in waking outcome measures for quantification of plasticity effects induced by the intervention. Across intervention sessions, the nocturnal naps began at 23:09 on average (SD, 60.83 min) and lasted 153.82 ± 77.15 min. Fast-spindle density during NREM sleep did not differ significantly among stimulation conditions (trough: 2.00 ± 1.17 spindles/min; peak: 2.00 ± 1.14; random-phase: 1.77 ± 0.71; spindle-free: 1.79 ± 1.70; p = 0.818). Mean fast-spindle duration also did not differ significantly among conditions (trough: 0.76 ± 0.06 s; peak: 0.78 ± 0.04 s random-phase: 0.75 ± 0.05 s; spindle-free: 0.76 ±0.05 s; p = 0.178). Among the spindle-triggered conditions, instantaneous spindle amplitude immediately before the TMS triplets was 12.26 ± 4.70 µV in the trough condition, 11.70 ± 4.49 µV in the peak condition, and 10.36 ± 3.03 µV in the random-phase condition (p = 0.068). Participants received 190.3 ± 44.5 spindle-peak, 178.9 ± 70.1 spindle-trough, 195.4 ± 19.2 random spindle-phase, and 202.0 ± 34.1 spindle-free EEG-triggered TMS triplets, with no significant difference in stimulation number across conditions (p = 0.577). The mean ITI differed significantly across conditions (p < .001), with SP-free condition (14.41 ± 14.06 s) shorter than in the peak (49.82 ± 21.73 s, p_corrected_ = .001), trough (51.28 ± 24.25 s, p_corrected_ = .003), and random conditions (37.17 ± 14.79 s, p_corrected_ = .004). Mean ITI was also shorter in the random condition than in the peak (p_corrected_ = .045) and trough conditions (p_corrected_ = .040), while peak and trough did not differ (p_corrected_ = .873).

### 3.1 Real-time EEG-triggered TMS targeting of sleep spindle states

Offline pre-trigger EEG analysis confirmed successful spindle-state targeting (Fig. 2). Circular statistics demonstrated non-uniform phase distributions for peak stimulation (Rayleigh test: z = 8.697, p < 0.001) and trough stimulation (Rayleigh test: z = 10.303, p < 0.001), with directed clustering toward the intended target phases (peak: V-test V = 0.748, p < 0.001; trough: V-test V = 0.838, p < 0.001). Random-phase stimulation showed no evidence of phase clustering (Rayleigh test: z = 0.407, p = 0.674).

### 3.2 Pre- and post-sleep analyses

No statistically significant between-condition differences were detected pre- intervention for any of the waking outcome measures (rs-EEG power, log10-transformed MEP amplitude, LMFA, TMS-induced EEG power, Table S1).

#### 3.2.1 Resting-state EEG power changes

Resting-state EEG high-beta power in the left sensorimotor ROI did not show significant differences between the four experimental conditions at Post10 (Fig. 3A). At Post30, the Pre-to-Post30 change in high-beta power was more negative after trough stimulation than after peak (estimate = -0.665 dB, 95% CI [-1.197, -0.132], p_corrected_ = 0.029), random (estimate = -0.844 dB, 95% CI [-1.387, -0.300], p_corrected_ = 0.007), and spindle-free stimulation (estimate = -1.135 dB, 95% CI [-1.668, -0.603], p_corrected_ < 0.001; Fig. 3B). Within the trough condition, high-beta power decreased at Post30 compared to Pre (estimate = −0.869 dB, 95% CI [−1.363, −0.374], p_corrected_ = 0.009; Fig. 3B). No other within-condition Post30 or Post10 comparisons with Pre were significant. No pairwise contrast in the mu/alpha, or low-beta band was significant after correction for multiple comparisons (all p_corrected_ > 0.05).

**Figure 3.**
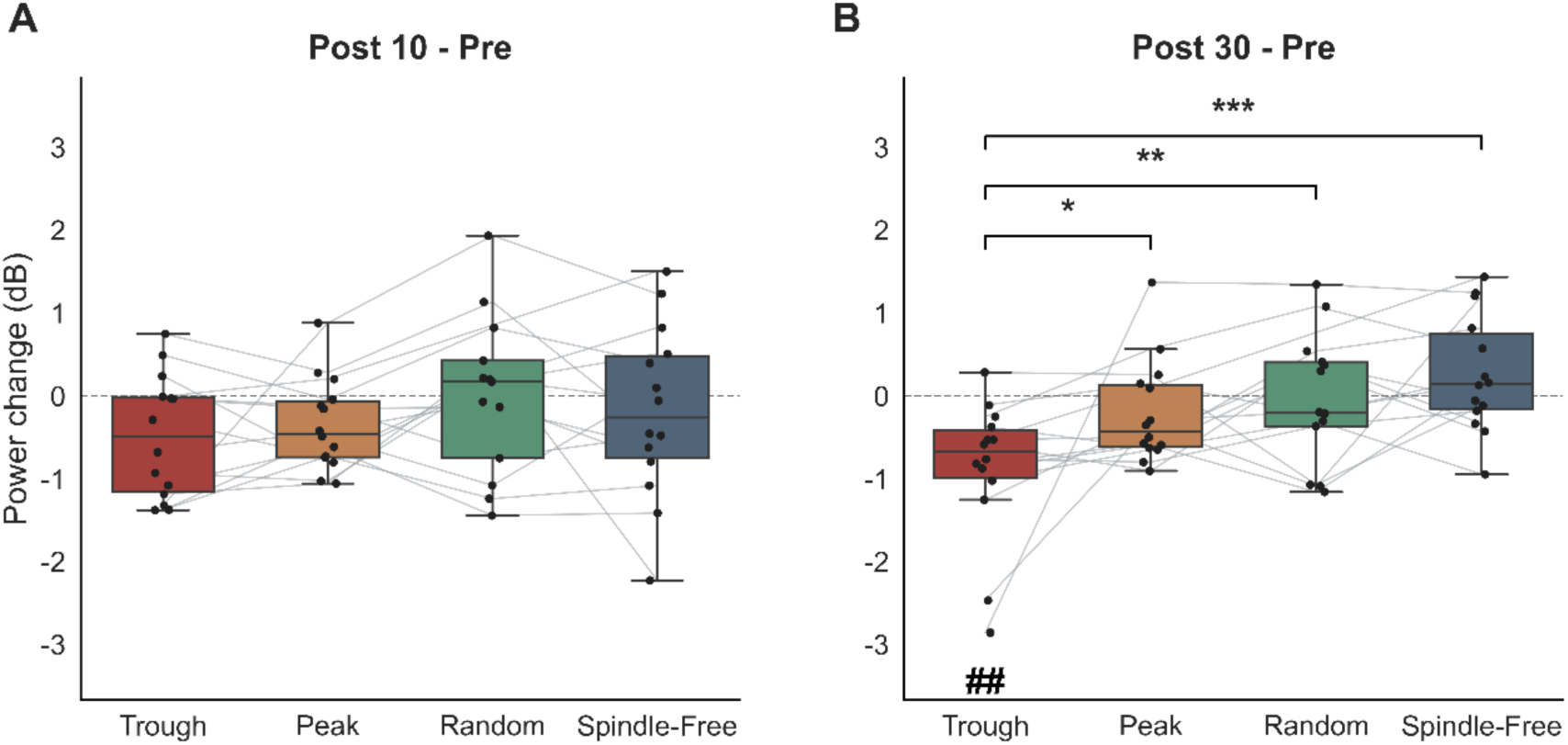
Changes in rsEEG high-beta power in the left sensorimotor ROI. (A) Post10-Pre and (B) Post30- Pre changes in high-beta (25-30 Hz) power across the four stimulation conditions. Boxes show the median and interquartile range; whiskers extend to the most extreme observation within 1.5 times the interquartile range. Dots represent the data of individual participants, and gray lines connect repeated measurements within participants. Brackets and asterisks indicate FDR-corrected significant pairwise condition contrasts (*p < 0.05, **p < 0.01, ***p < 0.001). Hash symbols below individual boxes indicate significant FDR-corrected Post-Pre changes within that condition (## p < 0.01).

#### 3.2.2 Changes in motor evoked potential (MEP) amplitude

Post-Pre changes in log10-transformed MEP amplitude did not differ significantly among conditions at Post10 (p_corrected_ = 0.123; Fig. 4A) or Post30 (p_corrected_ = 0.163; Fig. 4B). Analysis of Post-Pre changes within condition showed a significant Post10-Pre decrease after trough stimulation (estimate = −0.758 log10 µV, 95% CI [−1.196, −0.320], p_corrected_ = 0. 0099; Fig. 4A). No significant Post30-Pre change was detected within any condition.

**Figure 4.**
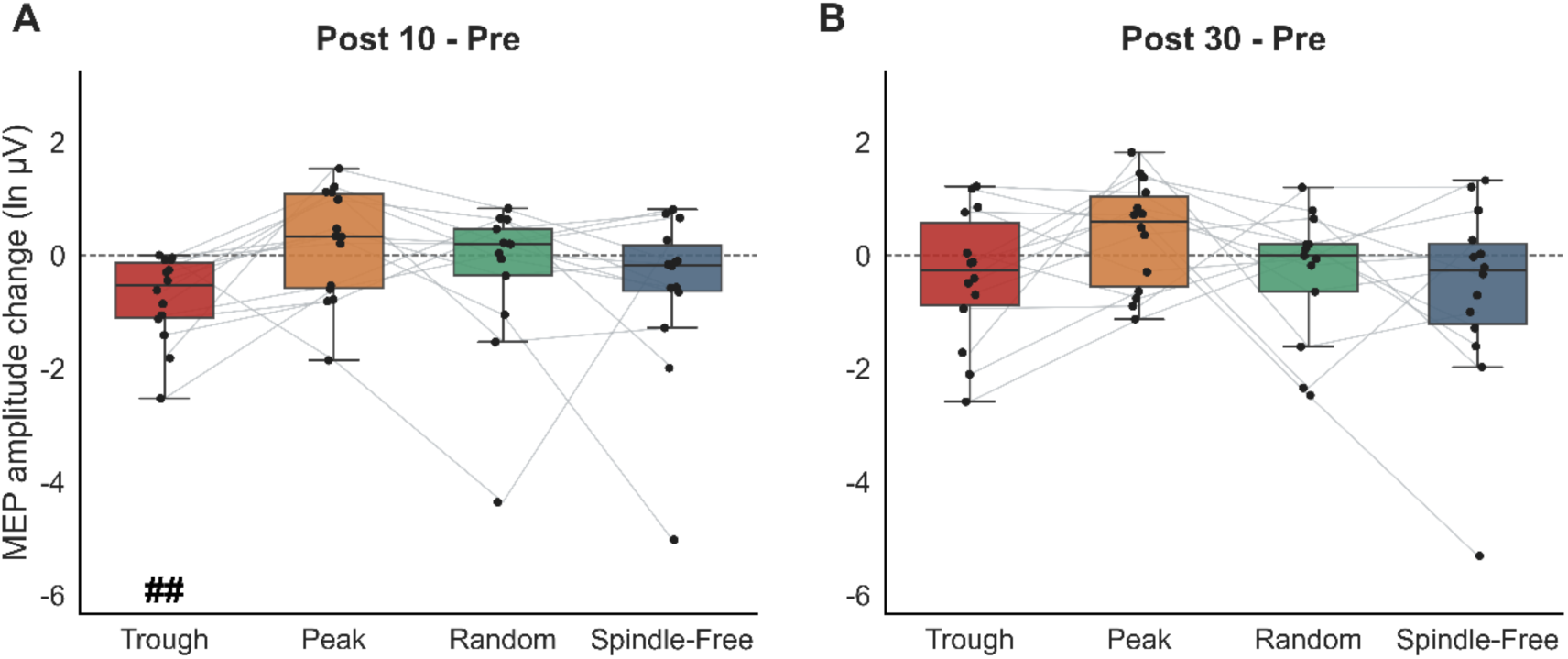
Changes in corticospinal excitability measured by log10 MEP amplitude. (A) Post10-Pre and (B) Post30-Pre changes across the four stimulation conditions. Boxes show the median and interquartile range; whiskers extend to the most extreme observation within 1.5 times the interquartile range. Dots represent the data of individual participants, and gray lines connect repeated measurements within participants. Hash symbols below individual boxes indicate significant FDR-corrected Post-Pre changes within that condition. ## p < 0.01.

#### 3.2.3 Changes in sensorimotor local mean field amplitude (LMFA)

Pairwise comparisons showed condition differences between experimental conditions in the N45 and P60 LMFA change scores (Fig. 5). At Post10, compared to pre-intervention, the P60 LMFA was more negative (i.e., the P60 became smaller) after trough stimulation than after peak (estimate = -0.77 μV, 95% CI [-1.31, -0.22], p_corrected_ = 0.018), random (estimate = -0.64 μV, 95% CI [-1.20, -0.08], p_corrected_ = 0.048), and spindle-free stimulation (estimate = -0.88 μV, 95% CI [-1.43, -0.33], p_corrected_ = 0.010; Fig. 5B).

**Figure 5.**
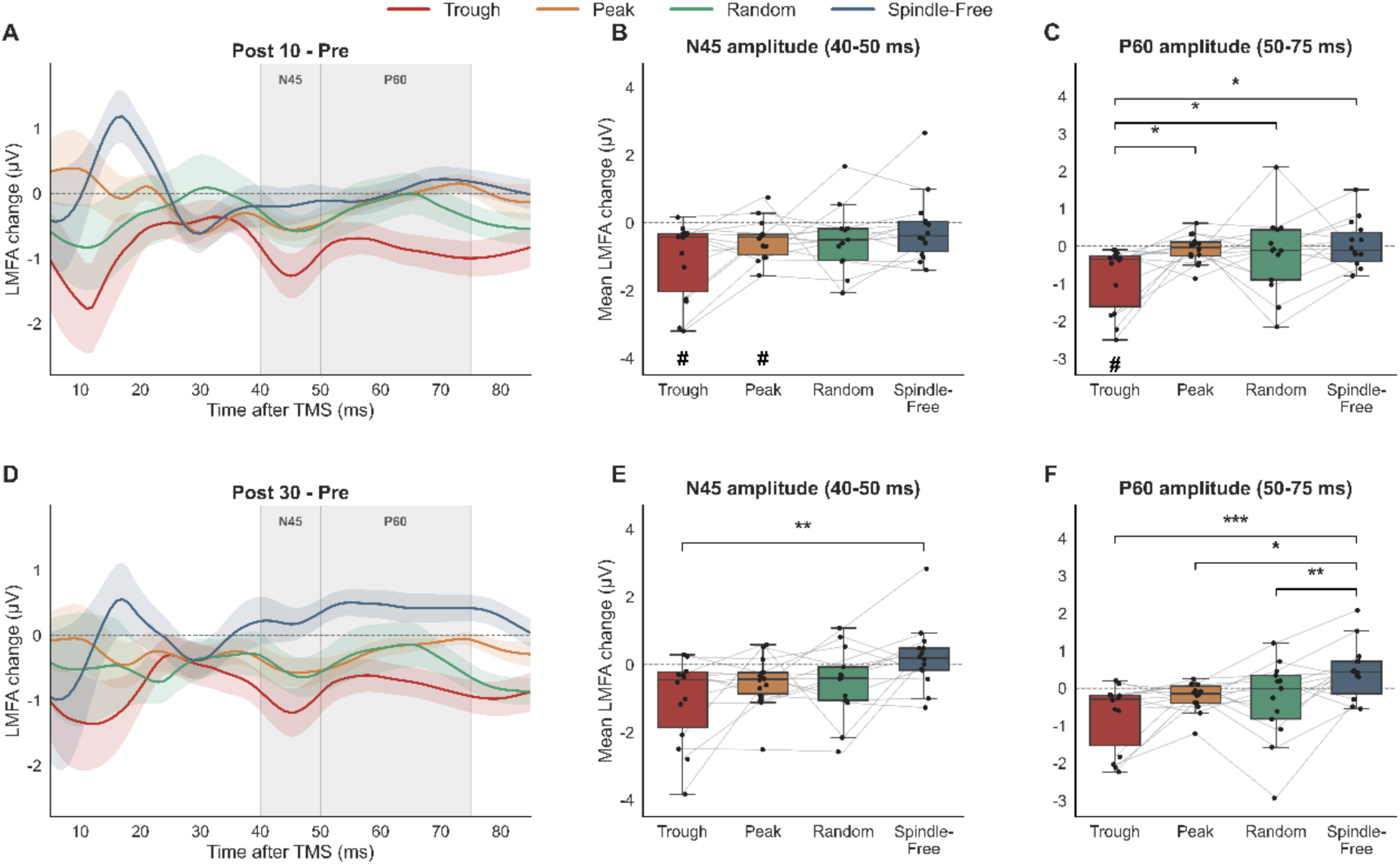
Changes in TMS-evoked local mean field amplitude (LMFA) in the left-sensorimotor ROI. (A) Group-level LMFA Post10-Pre changes. (B-C) Distributions of the mean N45 and P60 LMFA Post10-Pre changes, respectively. (D) Group-level LMFA Post30-Pre changes. (E, F) Distributions of the mean N45 and P60 LMFA Post30-Pre changes, respectively. In A and D, lines represent group means and shadings indicate ± SEM. Gray regions indicate the N45 (40-50 ms) and P60 (50-75 ms) analysis windows. In B-C, and E-F, dots represent data of individual participants, gray lines connect repeated within-participant measurements across experimental conditions, boxes show the median and interquartile range, and whiskers extend to 1.5 times the interquartile range. Brackets and asterisks indicate significant FDR-corrected pairwise differences between condition-specific change scores (*p < 0.05, **p < 0.01, ***p < 0.001). Hash symbols below individual boxes indicate significant FDR-corrected Post-Pre changes within that condition (# p < 0.05).

At Post30, the N45 LMFA change was more negative after trough than after spindle-free stimulation (estimate = -1.28 μV, 95% CI [-2.01, -0.55], p_corrected_ = 0.004; Fig. 5D). P60 LMFA changes were more negative after trough (estimate = -1.19 μV, 95% CI [-1.67, -0.70], p_corrected_ < 0.001), peak (estimate = -0.67 μV, 95% CI [-1.16, -0.18], p_corrected_ = 0.014), and random stimulation (estimate = -0.76 μV, 95% CI [-1.26, -0.26], p_corrected_ = 0.008) than after spindle-free stimulation (Fig. 5E) compared to pre-intervention. No other comparison between stimulation conditions was significant.

Analysis of Post-Pre within-condition changes showed significant Post10-Pre decreases in N45 LMFA in the trough condition (estimate = -1.089uV, 95% CI [-1.753, -0.425], p_corrected_ = 0.029; Fig. 5B) and peak condition (estimate = -0.499 µV, 95% CI [-0.843, - 0.154], p_corrected_ = 0.043; Fig. 5B), as well as in P60 LMFA in the trough condition (estimate = -0.837 µV, 95% CI [-1.336, -0.338], p_corrected_ = 0.029; Fig. 5C).

#### 3.2.4 Changes in immediate response slope (IRS)

No pairwise differences in IRS among the trough, peak and random-phase conditions were significant at either assessment. At Post10, IRS changes were more negative (i.e., less steep) in the trough, peak, and random-phase condition than the spindle-free condition (estimates = -0.213, -0.238, and -0.215 µV/ms; 95% CIs [-0.370, -0.056], [-0.395, -0.081], and [-0.376, -0.054], respectively; all p_corrected_ = 0.018). At Post30, IRS changes remained more negative in the trough and peak compared to the spindle-free condition (estimates = -0.252 and -0.235 µV/ms; 95% CIs [-0.411, -0.094] and [-0.393, -0.076], respectively; both p_corrected_ = 0.011), while the random vs spindle-free condition contrast was no longer significant (p_corrected_ = 0.214). In addition, IRS decreased Post30-Pre in the trough condition (p_corrected_ = 0.032, Fig. 6B).

**Figure 6.**
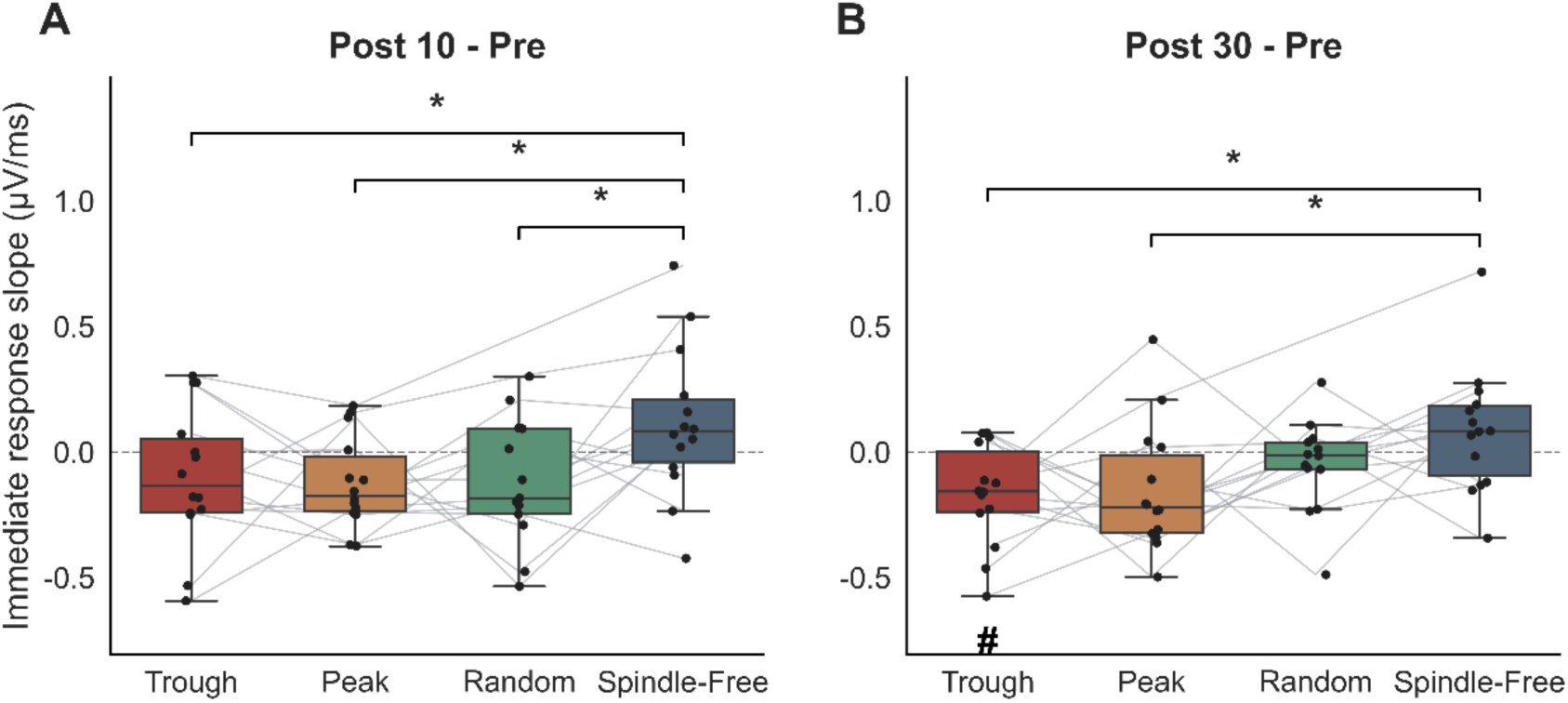
Changes in immediate response slope (IRS). (A) Post10-Pre and (B) Post30-Pre changes in IRS in the left sensorimotor ROI. Boxes show the median and interquartile range, whiskers extend to 1.5 times the interquartile range, dots represent the data of individual participants, and gray lines connect within-participant repeated measurements. Brackets indicate significant FDR-corrected contrasts between experimental conditions (*p < 0.05). The hash symbol below the trough box in panel B indicates a significant FDR-corrected Pre-Post30 change within the trough condition (# p < 0.05).

#### 3.2.5 Changes in time-frequency representations (TFRs)

The cluster-based comparison across the four conditions showed no significant differences in TFRs for Post10-Pre (p_corrected_ = 0.768) or Post30-Pre (p_corrected_ = 0.306). Analysis of within-condition Post-Pre changes of TFRs showed a significant negative Post30-Pre cluster in the trough condition, spanning 6-14 Hz and 0-300 ms after TMS (mean cluster change = -1.86 dB, p_corrected_ = 0.007; Fig. 7A-C). No significant Post30-Pre cluster was detected for peak, random, or spindle-free stimulation (Fig. 7D-F), nor for any condition Post10-Pre.

**Figure 7.**
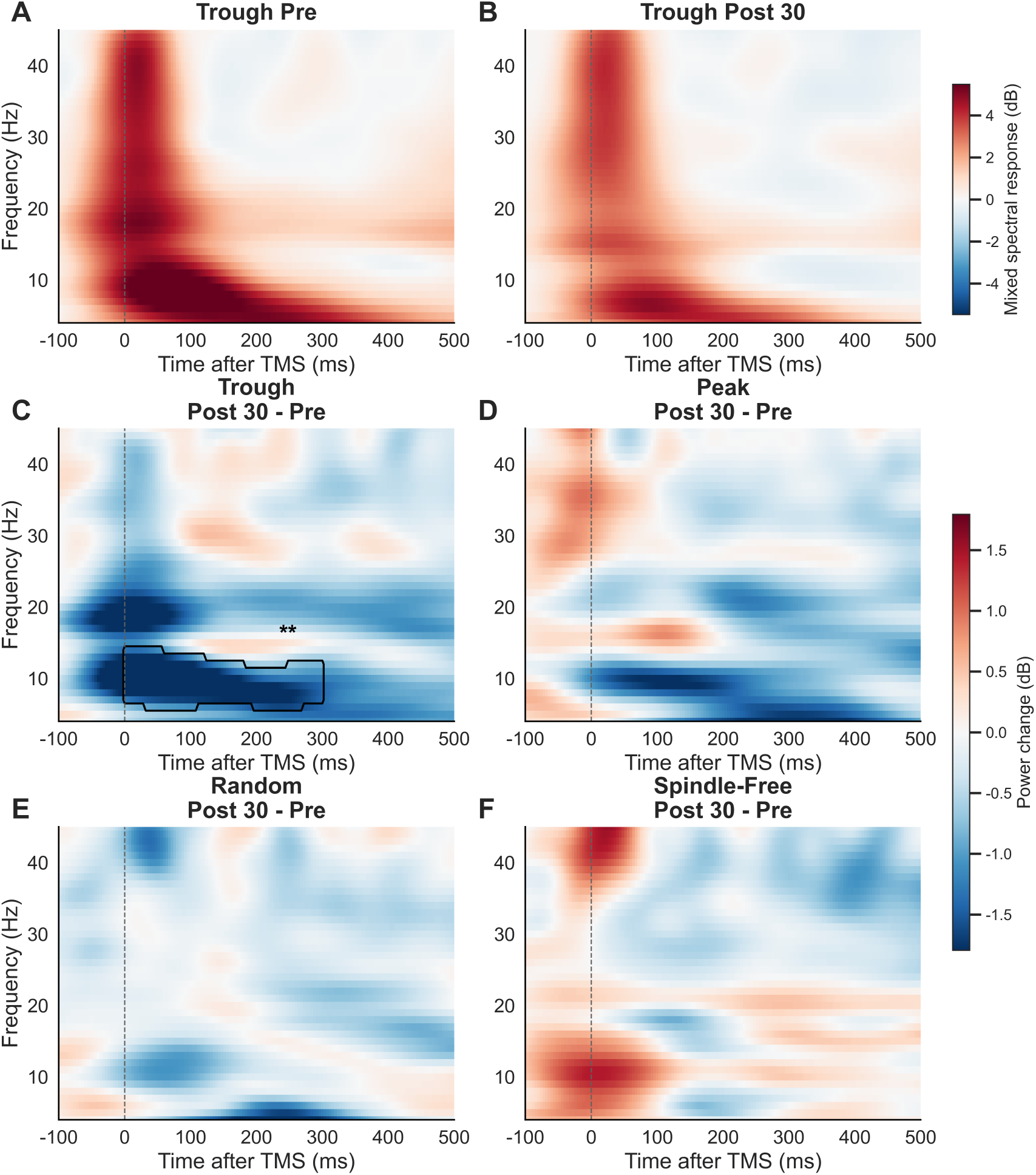
Changes in time-frequency representations (TFRs) in the left sensorimotor ROI. Group-average TFR of the TMS-related power change, containing evoked and induced responses at Pre (A) and Post30 (B) in the spindle-trough condition in the left sensorimotor ROI. Group-average Post30-Pre changes (in dB) in TFRs in the left sensorimotor ROI for the spindle-trough (C), spindle-peak (D), spindle random phase (E), and spindle-free condition (F). The vertical dashed line marks the time of the TMS pulse (0 ms). In panel C, the black contour marks the significant negative Post30-Pre TFR change in the spindle-trough condition within the prespecified 0-300 ms analysis window. Asterisks indicate FDR-corrected cluster-level significance across the four condition-specific tests. **p < 0.01.

## 4 Discussion

This study tested the effects of sleep spindle state-dependent burst TMS at hippocampal ripple frequency on plasticity induction in the human motor cortex. Across a variety of plasticity readouts (rs-EEG power, log10-transformed MEP amplitude, LMFA, IRS, TFR) the spindle-trough rTMS condition consistently stands out by leading to an LTD-like reduction of motor cortical excitability, significantly different from all other stimulation conditions (spindle-peak, random spindle phase, spindle-free). In the following paragraphs ewe provide a physiological interpretation of the observed changes in rsEEG, MEP and TMS-EEG metrics, followed by a discussion on possible explanations for why the changes were selectively induced by targeting the spindle-trough with ripple burst rTMS.

### 4.1 Changes in resting-state EEG power

Power in the beta-frequency band in the sensorimotor cortex in EEG, electrocorticography, and intracranial EEG recordings is desynchronized during voluntary movement execution (Crone et al., 1998; Stancák & Pfurtscheller, 1996; Szurhaj et al., 2003). This has generally been interpreted as an indicator of neuronal activity in the sensorimotor cortex (Pfurtscheller & Lopes da Silva, 1999), paralleled by increased MEP amplitude (Chen et al., 1998) and fMRI BOLD signal (Formaggio et al., 2008). In contrast, somatosensory evoked potential recordings show decreased responsiveness of the sensorimotor cortex during beta desynchronization (Cohen & Starr, 1987; Seki & Fetz, 2012), possibly an indication of decreased excitability. However, these studies relate to a voluntarily activated motor cortex, and the findings may not be directly applicable to a voluntarily relaxed state. In the resting state, high beta power is associated with high MEP amplitudes (Jin et al., 2026) and high early TEP amplitudes (Ahola et al., 2025; Perera et al., 2024). Therefore, the observed decrease in high-beta power specifically after spindle trough burst rTMS in this study (Fig. 3A-B) most likely represents a long-lasting excitability depression of the sensorimotor cortex. Similar positive correlations for rsEEG power in the mu-alpha frequency band with MEP amplitude (Ahola et al., 2025; Bergmann et al., 2019; Hussain et al., 2019; Ogata et al., 2019; Thies et al., 2018) and early TEP amplitude (Ahola et al., 2025; Perera et al., 2024) have been previously demonstrated. In the present study, however, spindle-trough burst rTMS specifically decreased rs-EEG power in the high-beta band, not in the alpha band. This specificity is compatible with observations from many other studies that sensorimotor alpha and beta oscillations are associated with distinct mechanisms and functions (for reviews, Kilavik et al., 2013; Pfurtscheller & Lopes da Silva, 1999).

### 4.2 Changes in motor evoked potential (MEP) amplitude

MEP amplitude is an extremely well established measure of corticospinal excitability (for reviews, Massimini et al., 2026; Siebner et al., 2022; Spampinato et al., 2023). Therefore, the observed decrease in MEP amplitude specifically by spindle-trough burst rTMS in this study (Fig. 4A) provides clear evidence for the induction of LTD-like plasticity of corticospinal excitability. A similar LTD-like change of MEP amplitude by brain state-dependent ∼1 Hz rTMS was previously demonstrated during wakefulness when specifically targeting the peak but not the trough of the ongoing sensorimotor mu-rhythm (Baur et al., 2020).

### 4.3 Changes in local mean field amplitude (LMFA)

Early TEP amplitudes in the sensorimotor cortex reflect excitability of the stimulated cortical networks (for reviews, Darmani & Ziemann, 2019; Ziemann et al., 2026). Pharmaco-TMS-EEG studies demonstrated that benzodiazepines, i.e., positive allosteric modulators at the GABAA receptor, and dextromethorphan, a non-competitive antagonist at the glutamatergic NMDA receptor, increase the N45 TEP amplitude (Belardinelli et al., 2021; Gordon et al., 2023; Premoli et al., 2014). Based on this background, the observed increase of the LMFA from EEG signal of the stimulated sensorimotor cortex in the N45 time window specifically by spindle-trough burst rTMS (Figs. 5A-B, D-E) can be interpreted as a decrease in excitability, either driven by an increase in GABAAergic inhibitory and/or a decrease in glutamatergic excitatory neurotransmission.

The P60 TEP amplitude is decreased by perampanel, a specific antagonist at ionotropic glutamatergic AMPA receptors (Belardinelli et al., 2021). The observed decrease of the LMFA in the P60 time window in this study specifically by spindle-trough burst rTMS (Fig. 5A, C, D, F) is therefore best interpreted as a decrease in cortical excitability related to fast ionotropic glutamatergic neurotransmission.

### 4.4 Changes in the immediate response slope (IRS)

Changes in the IRS, i.e., the ascending slope towards the P30 TEP is considered a marker of synaptic plasticity, with increases indicating LTP and decreases LTD (Casarotto et al., 2013; Massimini et al., 2026; Vyazovskiy et al., 2008). Electroconvulsive therapy in patients with treatment-refractory major depressive disorder resulted in concordant improvement of depression, increase in the P30 TEP amplitude and an increase in the IRS (Casarotto et al., 2013). Here we observed a decrease after all spindle-triggered burst rTMS protocols (Fig. 6) compared to stimulation during spindle-free periods. While this finding was not specific to the spindle-trough condition it corroborates the other findings of this study that spindle-trough burst rTMS leads to LTD-like corticospinal excitability.

### 4.5 Changes in time-frequency representations (TFRs)

The physiological underpinnings of TFRs are less well understood compared to the other metrics analyzed in this study (for reviews, Q. Wang et al., 2024; W. Wang et al., 2026; Ziemann et al., 2026). Here, we obtained the power of the TFR, also called TMS-related spectral perturbation. This is an extension of event-related spectral perturbation to TMS (Makeig, 1993). It measures the average dynamic change in evoked and induced oscillatory power after the TMS pulse across all the bands of the EEG frequency spectrum (Ziemann et al., 2026). TRFs have been related to the excitation/inhibition balance in cortical networks. XEN1101, a novel anti-seizure drug, which potentiates the open state of KCNQ2/3 potassium channels and thereby suppresses neuronal excitability, deceased mixed TFR oscillatory power in the alpha-band (Biondi et al., 2022).

The current findings show that burst rTMS specifically at the spindle-trough decreases TMS-related oscillatory power in the theta/alpha-frequency bands (Fig. 7A-C), consistent with the notion that this intervention is associated with a reduction of neural excitability in the stimulated sensorimotor cortex.

### 4.6 Why were the observed changes specific to the spindle-trough burst rTMS condition?

In previous studies, we and others have demonstrated that corticospinal excitability during wakefulness depends on the phase of the ongoing sensorimotor mu-rhythm. MEP amplitudes are largest at the trough and the ascending phase, indicating a relative corticospinal high-excitability state, and lowest during the peak and descending phase, indicating a relative corticospinal low-excitability state (Bergmann et al., 2019; Schaworonkow et al., 2018, 2019; Wischnewski et al., 2022; Zrenner et al., 2018, 2023). According to the principle of cooperativity in synaptic plasticity that predicts induction of LTP or LTD with depolarization vs. hyperpolarization of postsynaptic target cells (Sjöström et al., 2001), and in line with seminal in-vitro work in hippocampal CA1 neurons (Huerta & Lisman, 1995), we showed in real-time EEG-TMS experiments in humans that rTMS repeatedly and consistently targeting the high-excitability state (i.e., the trough of the ongoing mu-rhythm) results preferably in an LTP-like increase in MEP amplitude (Baur et al., 2022; Zrenner et al., 2018), while repeatedly targeting the low- excitability state (i.e., the peak of the ongoing mu-rhythm) leads preferably to LTD-like plasticity (Baur et al., 2020).

The oscillatory state-dependent corticospinal excitability during sleep is less well explored. It was demonstrated earlier that depolarized SO up-states vs. hyperpolarized SO down-states result in relatively larger vs. smaller MEPs, respectively, even gradually depending on the oscillatory amplitude at the stimulation site (Bergmann et al., 2012), but no comparison to desynchronized NREM baseline was made, and also phase-dependent plasticity by repeated stimulation of the same SO phase has not been investigated yet. More recently, we showed that single-pulse TMS during sleep spindles result in decreased MEP amplitude compared to TMS during spindle-free periods, with the falling flank of the spindle being the phase associated with lowest corticospinal excitability (Hassan et al., 2025), findings that we now largely replicated in an independent study (Breuer et al., unpublished data). Notably, no spindle phase was associated with corticospinal excitability increases relative to spindle-free baseline NREM sleep, suggesting spindles to be events of asymmetric ‘pulsed inhibition’ (Hassan et al., 2025), in line with the spindle-associated GABAergic activity observed in mice (Niethard et al., 2018), even though no phasic modulation of GABAergic paired-pulse short-interval intracortical inhibition could be observed in humans (Breuer et al., unpublished data). Therefore, our finding that burst rTMS in all spindle-locked conditions, but in particular when targeting the spindle-trough, resulted in LTD-like excitability decreases, may be similarly consistent with the principle of cooperativity in synaptic plasticity induction. While the oscillatory trough seems to be a phase particularly susceptible to fast input activity inducing synaptic changes in both wakefulness and sleep, the direction of change towards LTP- or LTD-like excitability modulation appears to rather depend on the direction of the asymmetric net excitability modulation associated with the temporally surrounding oscillatory event (i.e., facilitation during wake mu-alpha and inhibition during sleep spindles). While this interpretation is speculative, it raises interesting questions regarding the direction of phase-dependent plasticity effects that would result from exclusively targeting (inhibitory) spindles occurring during (facilitatory) depolarized SO up-states, to be investigated in future studies.

Moreover, the trough is the natural spindle phase in which hippocampal ripples are predominantly nested (Staresina et al., 2015). Active systems memory consolidation critically depends on the hierarchical interaction of sleep oscillations across the hippocampo-thalamo-cortical system (Klinzing et al., 2019; Lutz et al., 2026; Staresina, 2024; Staresina et al., 2015): Neocortical slow oscillations (<1 Hz) synchronize cortical networks by alternating depolarizing up-states and hyperpolarizing down-states, temporally grouping thalamo-cortical sleep spindles (12–15 Hz) into up-states, which in turn coordinate hippocampal sharp-wave ripples (>80 Hz), and thus the replay of memory traces. Slow oscillation-spindle-ripple events are therefore probably providing a critical window of enhanced synaptic plasticity across large-scale networks responsible for memory consolidation (Bergmann & Born, 2018; Klinzing et al., 2019; Staresina, 2024).

In the mouse hippocampus, sharp-wave ripple oscillations serve as intrinsic events that trigger LTD, presumably to erase non-relevant memory traces (Norimoto et al., 2018). Therefore, injecting ripple bursts exogenously into the spindle trough may support the intrinsic process of pruning memory traces through LTD-like plasticity. The trough-specific induction of LTD-like plasticity demonstrated in this study opens the intriguing possibility of targeted manipulation of spindle-ripple coupling for impacting on human behaviour, such as motor memory consolidation. If indeed the specific excitability profile temporally surrounding the stimulation event is determining the direction of plasticity induction, as speculated above, it will be important to experimentally disentangle the respective contributions of the event- and phase-related excitability modulations associated with SOs, spindles, and their specific temporal interaction, the latter believed to establish a particularly relevant state for synaptic rescaling during sleep through the differential activation of different inhibitory interneuron subclasses (Klinzing et al., 2019; Niethard et al., 2018; Rolle et al., 2025). We plan to examine this in subsequent studies, directly building on the here presented findings.

### 4.7 Limitations

This study has several limitations. First, the final sample was relatively small but in accord with our sample size estimation. Still, reliability of effect estimates is limited and generalizability will need to be validated in independent larger participant cohorts. Second, the stimulation conditions also covered only selected points of the spindle cycle. We targeted sleep spindle peak, trough, and random-phase, but did not include rising- or falling-phase stimulation. This may be potentially of interest in future experiments as there is evidence that it is the falling phase that his associated with the strongest corticospinal excitability decrease (Hassan et al., 2025). Third, rTMS targeting of spindles did not distinguish between spindles nested in slow oscillations (SOs) vs. isolated spindles outside of slow oscillations. The principal condition effects remained after including SO exposure and spindle-SO nesting in the models (see Supplementary Materials, Tables S2-S3), suggesting that they were not readily explained by differences in the proportion of rTMS bursts delivered during SOs or the proportion of stimulated spindles that were SO-nested. However, these exploratory analyses do not establish that the observed effects were independent of SO physiology. Our interventions did not separately target SO-nested vs. isolated spindles or specific SO phases, and the exploratory interaction analyses were limited by the overlap in SO exposure across conditions. The present analyses therefore remain consistent with a spindle-trough effect while leaving open whether precise spindle-SO coupling further modulates the magnitude of the observed aftereffects. This likely is a particularly important distinction as the hippocampo-thalamo-cortical trialogue with nesting of ripples in the trough of spindles and nesting, in turn, of spindles in the up-state of slow oscillations is thought to be the electrophysiological hallmark of plasticity induction underlying memory consolidation (Bergmann & Born, 2018; Klinzing et al., 2019; Staresina, 2024). We will experimentally address this question in upcoming experiments utilizing novel real-time EEG-TMS software that will enable the reliable online distinction of SO-nested vs. free spindles (Kahilakoski et al., 2025). Fourth, the post-nocturnal nap measures may have been affected non-specifically by residual sleepiness differences. Participants were kept awake and allowed to move briefly after awakening, but sleepiness or alertness were not quantified. Finally, we have not tested effects of burst rTMS on behavior, in particular memory consolidation. We plan to examine this in subsequent studies.

## 5 Conclusions

This is first realization of real-time EEG-ripple burst rTMS to target specific phases of ongoing sleep spindles in human cortex for plasticity induction. Findings demonstrate that targeting the spindle trough stands out from all other experimental conditions (spindle-peak, spindle random phase, spindle-free) in showing consistent features of LTD-like depression across all readouts of corticospinal and sensorimotor cortical excitability. This opens the intriguing opportunity of targeted manipulation of human sleep physiology using non-invasive brain stimulation (Krugliakova et al., 2026) for improving specific behavioral processes, such as memory consolidation.

## Acknowledgements

We thank Dr. Umair Hassan for development and implementation of the software that has been used here for real-time EEG sleep spindle detection.

## CRediT author contributions

**Zhijian Zhao:** Writing – review & editing, Writing – original draft, Visualization, Validation, Software, Resources, Project administration, Methodology, Investigation, Formal analysis, Data curation, Conceptualization.

**Yeyun Lu:** Writing – review & editing, Writing – original draft, Visualization, Validation, Software, Resources, Project administration, Methodology, Investigation, Formal analysis, Data curation, Conceptualization.

**Friederike Breuer:** Writing – review & editing, Methodology, Conceptualization.

**Til Ole Bergmann:** Writing – review & editing, Supervision, Funding acquisition, Formal analysis, Conceptualization.

**Ulf Ziemann:** Writing – review & editing, Supervision, Software, Resources, Project administration, Funding acquisition, Formal analysis, Data curation, Conceptualization.

## Data and materials availability

The data and materials that support the findings of this study are available from the corresponding author upon reasonable request.

## Supplementary Materials

### Resting-state EEG quality control

Bad channels were removed before ICA and subsequently reconstructed by spherical interpolation. Across the 165 rsEEG recordings included in the analysis, 2.71 ± 2.07 channels per recording were interpolated (median, 2; range, 0-9), and 6.14 ± 3.10 components per recording were removed by ICA (median, 6; range, 0-16).

### Slow oscillation (SO) context of spindle-triggered TMS trials

The SO-spindle-ripple events are thought to be critical for plasticity induction and memory consolidation (Bergmann & Born, 2018; Klinzing et al., 2019; Staresina, 2024). Although we have not specifically tested burst rTMS interventions targeting specifically SO-nested spindles vs. free spindles in separate experimental sessions, there was inter-individual variation of the proportion of these events.

To assess whether differences in SO context could contribute to the plasticity effects of spindle-triggered burst rTMS, we quantified the proportion of spindle-targeted trials occurring within a detected SO. SO-nested trials were further classified according to whether stimulation occurred during the SO up-state or down-state.

The proportion of SO-nested stimulation trials was 43.81 ± 6.72% in the trough condition, 41.27 ± 11.70% in the peak condition, and 45.05 ± 11.98% in the random-phase condition. Nesting proportions did not differ among the three spindle-triggered conditions (p = 0.584). Among SO-nested trials, stimulation occurred predominantly during the SO up-state, comprising 80.26%, 81.85%, and 85.31% of nested trials in the trough, peak, and random-phase conditions, respectively. The corresponding SO down-state proportions were 19.74%, 18.15%, and 14.69%, respectively. The up/down-state distribution of SO-nested trials did not differ among conditions (p = 0.292). We additionally quantified SO exposure, defined as the proportion of trials with stimulation occurring within a detected SO, across all four intervention conditions. SO exposure differed significantly across the four intervention conditions (p < 0.001). Exposure was lower in the spindle-free condition (17.23 ± 14.86%) than in the trough (p = 0.002), peak (p < 0.001), and random-phase conditions (p = 0.001).

### Sensitivity analyses of SO context and spindle-SO nesting

To examine to what extent SO context contributed to variability in the Post-Pre corticospinal and sensorimotor cortical excitability changes, Post10-Pre and Post30-Pre changes were analyzed separately using linear mixed-effects models. SO-related predictors were centered within participant and scaled in 10-percentage-point units. SO exposure was the percentage of TMS bursts occurring during a detected SO. The SO phase was centered within participant. Positive values indicate relatively greater exposure during the SO up-state, and negative values indicate greater exposure during the SO down-state. The model across all four intervention conditions (spindle-trough, spindle-peak, spindle-random-phase, and spindle-free stimulation) included condition, SO exposure, and SO phase, and ΔY denotes the corresponding Post10-Pre or Post30-Pre change:

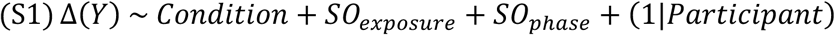

For the three spindle-targeted conditions, changes were modeled as a function of condition, spindle-SO nesting, and SO phase quantified within SO-nested trials:

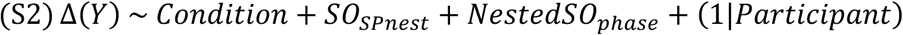

Here, SO_SPnest_ indicates the percentage of stimulated spindles that were nested within an SO. Nested SO phase describes whether SO-nested stimulated spindles occurred more frequently during the SO up- or down-state.

An exploratory model additionally tested whether the association between SO exposure and outcome change differed across intervention conditions:

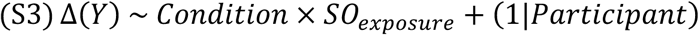

Neither SO exposure nor SO phase was significantly associated with changes in high-beta rsEEG power, MEP amplitude, N45 or P60 LMFA, or IRS after FDR correction (all p_corrected_ ≥ 0.838; Table S2). For the three spindle-targeted conditions, neither spindle-SO nesting nor SO up-down balance within nested trials was significantly associated with any outcome (all p_corrected_ ≥ 0.885; Table S3). There was also no evidence that the association between SO exposure and outcome change differed across intervention conditions (all p_corrected_ ≥ 0.327).

---

**Table S1.** Pre-intervention waking outcome measures across experimental conditions. (spindle-trough, spindle-peak, spindle random phase, spindle-free). Values are means ± SD. P values are from omnibus condition tests in linear mixed-effects models with participant-specific random intercepts, and were FDR-corrected across variables within each outcome family. Note that all p-values are > 0.05.

| Outcome family | Variable | Unit | Trough | Peak | Random | Spindle-Free | p<sub>corrected</sub> |
| --- | --- | --- | --- | --- | --- | --- | --- |
| rsEEG power | mu/alpha | dB | 4.32 ± 3.57 | 2.92 ± 3.23 | 3.81 ± 3.46 | 3.51 ± 3.92 | 0.081 |
| | low-beta | dB | 1.95 ± 1.63 | 1.36 ± 1.83 | 1.96 ± 1.91 | 1.63 ± 1.88 | 0.118 |
| | high-beta | dB | 1.56 ± 1.39 | 1.00 ± 1.14 | 1.12 ± 1.25 | 0.95 ± 1.32 | 0.081 |
| MEP amplitude | Log10-transformed MEP | log10 μV | 2.47 ± 0.57 | 2.47 ± 0.44 | 2.50 ± 0.41 | 2.49 ± 0.49 | 0.990 |
| Mean LMFA | N15/early | μV | 3.13 ± 1.28 | 2.63 ± 1.44 | 3.10 ± 1.50 | 2.65 ± 1.65 | 0.779 |
| | P30 LMFA | μV | 1.84 ± 1.22 | 1.89 ± 0.98 | 1.82 ± 1.26 | 1.83 ± 0.83 | 0.990 |
| | N45 | μV | 2.51 ± 1.93 | 1.84 ± 1.03 | 2.00 ± 0.87 | 1.78 ± 0.84 | 0.164 |
| | P60 | μV | 1.94 ± 1.27 | 1.39 ± 0.68 | 1.72 ± 0.99 | 1.32 ± 0.64 | 0.148 |
| Mean IRS | Mean IRS | μV/ms | 0.41 ± 0.31 | 0.41 ± 0.35 | 0.36 ± 0.34 | 0.26 ± 0.22 | 0.236 |
| TFR | mu/alpha | dB | 4.28 ± 1.75 | 3.80 ± 2.25 | 3.22 ± 1.46 | 3.21 ± 1.86 | 0.104 |
| | low-beta | dB | 2.30 ± 1.05 | 1.98 ± 1.39 | 1.77 ± 1.58 | 1.78 ± 1.12 | 0.693 |
| | high-beta | dB | 1.29 ± 0.91 | 1.09 ± 1.64 | 0.90 ± 1.63 | 1.19 ± 1.51 | 0.875 |

**Table S2.** Associations of slow-oscillation context with post-intervention changes across the four intervention conditions. Coefficients represent the change in the outcome per 10-percentage-point within-participant increase in the corresponding SO-related predictor. Values within cells are listed as Post10-Pre followed by Post30-Pre. Models included spindle-trough, spindle-peak, spindle random-phase, and spindle-free. FDR correction was applied within the prespecified analysis families.

| Outcome (unit) | Assessment | SO exposure<br>$\beta$ [95% CI] | $p_{\text{corrected}}$ | SO up-down balance<br>$\beta$ [95% CI] | $p_{\text{corrected}}$ | Condition × SO<br>$\chi^2(3)$ | $p_{\text{corrected}}$ |
| --- | --- | --- | --- | --- | --- | --- | --- |
| High-beta rsEEG (25-30 Hz), dB | Post10-Pre | -0.106 [-0.353, 0.141] | 0.838 | 0.071 [-0.220, 0.362] | 0.999 | 1.119 | 0.955 |
|  | Post30-Pre | -0.095 [-0.372, 0.182] | 0.838 | 0.078 [-0.248, 0.404] | 0.999 | 0.256 | 0.968 |
| MEP amplitude, log10(μV) | Post10-Pre | -0.085 [-0.210, 0.039] | 0.838 | 0.081 [-0.065, 0.228] | 0.999 | 8.760 | 0.327 |
|  | Post30-Pre | -0.048 [-0.187, 0.092] | 0.838 | 0.000 [-0.164, 0.164] | 0.999 | 6.013 | 0.370 |
| N45 LMFA, μV | Post10-Pre | -0.136 [-0.485, 0.214] | 0.838 | -0.040 [-0.451, 0.371] | 0.999 | 2.703 | 0.925 |
|  | Post30-Pre | 0.008 [-0.373, 0.388] | 0.985 | -0.047 [-0.494, 0.401] | 0.999 | 6.598 | 0.370 |
| P60 LMFA, μV | Post10-Pre | -0.162 [-0.442, 0.118] | 0.838 | 0.059 [-0.270, 0.389] | 0.999 | 2.085 | 0.925 |
|  | Post30-Pre | -0.031 [-0.286, 0.223] | 0.985 | 0.065 [-0.235, 0.364] | 0.999 | 0.848 | 0.955 |
| IRS, μV/ms | Post10-Pre | 0.001 [-0.082, 0.083] | 0.985 | -0.005 [-0.102, 0.092] | 0.999 | 0.759 | 0.955 |
|  | Post30-Pre | 0.020 [-0.060, 0.099] | 0.900 | -0.067 [-0.161, 0.026] | 0.999 | 2.345 | 0.925 |

**Table S3.** Associations of spindle-slow-oscillation nesting with post-intervention changes across the three spindle-targeted conditions. Coefficients represent the change in the outcome per 10-percentage-point within-participant increase in the corresponding SO-related predictor. Values within cells are listed as Post10-Pre followed by Post30-Pre. Models included spindle-trough, spindle-peak, and spindle random-phase. FDR correction was applied within the prespecified analysis families.

| Outcome (unit) | Assessment | Spindle-SO nesting<br>$\beta$ [95% CI] | $p_{\text{corrected}}$ | Nested SO up-down balance<br>$\beta$ [95% CI] | $p_{\text{corrected}}$ |
| --- | --- | --- | --- | --- | --- |
| High-beta rsEEG (25-30 Hz), dB | Post10-Pre | 0.032 [-0.307, 0.371] | 0.913 | -0.056 [-0.438, 0.327] | 0.947 |
|  | Post30-Pre | 0.042 [-0.373, 0.458] | 0.913 | -0.147 [-0.616, 0.322] | 0.947 |
| MEP amplitude, log10(μV) | Post10-Pre | 0.052 [-0.103, 0.208] | 0.913 | -0.050 [-0.225, 0.126] | 0.947 |
|  | Post30-Pre | 0.141 [-0.040, 0.322] | 0.913 | -0.141 [-0.345, 0.064] | 0.885 |
| N45 LMFA, μV | Post10-Pre | 0.028 [-0.478, 0.534] | 0.913 | -0.126 [-0.696, 0.444] | 0.947 |
|  | Post30-Pre | 0.161 [-0.368, 0.690] | 0.913 | -0.020 [-0.617, 0.577] | 0.947 |
| P60 LMFA, μV | Post10-Pre | -0.104 [-0.541, 0.333] | 0.913 | -0.024 [-0.517, 0.468] | 0.947 |
|  | Post30-Pre | -0.043 [-0.436, 0.351] | 0.913 | 0.017 [-0.428, 0.461] | 0.947 |
| IRS, μV/ms | Post10-Pre | -0.030 [-0.152, 0.093] | 0.913 | 0.060 [-0.078, 0.199] | 0.947 |
|  | Post30-Pre | 0.078 [-0.037, 0.192] | 0.913 | -0.092 [-0.222, 0.037] | 0.885 |

